# Development of approaches to overcome the drop in hematocrit when implementing mononuclear phagocyte system cytoblockade *in vivo* used to prolong the circulation of nanoparticles in the blood

**DOI:** 10.64898/2026.05.04.722692

**Authors:** Elizaveta N. Mochalova, Maria A. Yurchenko, Marina P. Timofeeva, Darina A. Maedi, Petr I. Nikitin, Maxim P. Nikitin

## Abstract

While engineered nanomaterials offer unprecedented precision in targeting tumor cells, their efficacy is often limited by rapid clearance from the bloodstream via the mononuclear phagocyte system (MPS). To overcome this limitation, a promising strategy known as MPS-cytoblockade has been developed. This approach involves administering antibodies against host erythrocytes. The resulting saturation of the MPS with erythrocyte clearance creates a critical window, allowing subsequently administered nanoparticles to evade immune surveillance and circulate for a significantly extended period. However, MPS-cytoblockade induces a transient reduction in hematocrit, which can lead to adverse effects. Here, we demonstrate that approaches to restore hematocrit, specifically through the administration of donor erythrocyte suspension or the hormone erythropoietin, effectively prevent this drop while maintaining the efficacy of the MPS-cytoblockade. Notably, these interventions do not compromise the prolonged circulation time of the nanoparticles or alter their biodistribution, preserving high accumulation in tumors. Our findings establish a viable strategy to mitigate a key side effect of MPS-cytoblockade, thereby enhancing its therapeutic potential and safety profile.

## 1. Introduction

Nanomedicine emerged as a strategy to overcome the limitations of traditional methods for disease diagnosis and treatment. It involves the development and use of materials at the nanoscale to create vaccines, biosensors, transfection systems, as well as new bioimaging tools. Another key area of nanomedicine is the creation of targeted drug delivery systems based on nanoparticles (NPs), which are used primarily for the therapy of oncological diseases [1], [2], [3], [4]. The approach reduces the toxic effect of chemotherapeutic drugs on normal, non-pathogenic cells by ensuring targeted action only on tumor tissues. Targeting can be achieved through the accumulation of NPs in the permeable vasculature endothelium of tumor tissue (the so-called EPR effect) or through the use of recognizing bioligands (e.g., antibodies) for specific binding to the target.

However, NPs entering the bloodstream are rapidly captured by the mononuclear phagocyte system (MPS) and cleared from circulation. As a result, the NPs accumulate primarily in the liver and spleen, failing to reach target organs and tissues [5]. This process constitutes a key obstacle to the broader clinical adoption of NPs. Several strategies have been previously proposed to reduce the uptake of nanomedicines by the MPS and thereby prolong their circulation time in the blood. Some involve modifying NPs with various coatings to achieve stealth properties, for example, using polyethylene glycol (PEG) [6] [7], while others aim to reduce the phagocytic activity of MPS cells by saturating them with various blocking agents (e.g., macrophage-toxic liposomal clodronate [8] [9] or high doses of “decoy” NPs [10] [11]).

A promising MPS-cytoblockade method was recently proposed [1], based on the administration of anti-erythrocyte antibodies. These antibodies “mark” red blood cells for phagocytosis by MPS cells, thereby inducing their saturation. Upon subsequent injection of NPs, they become “invisible” to the body’s immune system. This significantly prolongs their circulation time and, consequently, substantially increases the efficiency of their accumulation in target organs or tissue. Specifically, this approach allowed to increase the circulation half-life of a range of nanoparticle formulations by up to 32-fold. Importantly, the method does not require modification of the nanoparticle surface itself, which underlies the applicability of this technology to a wide range of nanoparticles of various materials and sizes, as well as to nanoagents functionalized with target-recognizing bioligands or complex, so-called “smart,” interfaces for analyzing the cellular microenvironment [12]. MPS-cytoblockade has great potential for safe use, as it accelerates the natural process of clearing the body’s own red blood cells without compromising its protective functions [13]. Furthermore, the technology has a high probability of successful translation into clinical practice, as anti-erythrocyte antibodies are currently approved for use in the treatment of immune thrombocytopenia [14].

Nevertheless, application of MPS-cytoblockade is associated with a temporary decrease in hematocrit levels, which may cause undesirable side effects. Maintaining normal hematocrit is particularly crucial because cancer patients often already suffer from anemia due to various factors. Anemia may be associated with increased expression of inflammatory cytokines that reduce red blood cell survival, impaired iron metabolism, insufficient production of the hormone erythropoietin, as well as the use of cytostatic drugs during chemotherapy, which leads to suppression of erythroid progenitor cells [15].

This study aims to develop approaches for restoring hematocrit levels during MPS-cytoblockade, specifically through the administration of donor blood components and the hormone erythropoietin, which stimulates red blood cell production in the bone marrow. We examined how these interventions affected nanoparticle circulation, biodistribution, and the efficiency of accumulation in the tumor. The developed strategies could help overcome the main side effect of MPS-cytoblockade and subsequently improve the performance and safety of a broad spectrum of nanomedicines for the treatment of oncological and other socially significant diseases.

## 2. Results and Discussion

### 2.1. The impact of MPS-cytoblockade on nanoparticle circulation time, tumor accumulation efficiency, and hematocrit level

First, we validated the potential of the MPS-cytoblockade method to increase both the circulation time of nanomaterials in the bloodstream and their accumulation in target tissues, particularly tumors.

The circulation kinetics of nanoparticles in the bloodstream of mice were studied *in vivo* using the highly sensitive magnetic particle quantification (MPQ) method, which enables the real-time quantification of nonlinear magnetic materials [1] [16]. To perform the measurements, the tail of an anesthetized mouse was placed in the detection coil of the device, followed by the retro-orbital injection of 100 nm glucuronic acid-coated fluidMAG-ARA nanoparticles (Chemicell, Germany), which served as model magnetic nanoparticles. To induce MPS-cytoblockade, antibodies such as mouse anti-mouse red blood cell clone 34-3C (Hycult Biotech, Netherlands) or rat anti-mouse red blood cell clone TER-119 (BioXCell, USA) at a dose of 1.25 mg/kg (25 µg per mouse) were injected 12 hours prior to NPs injection as demonstrated in the original study [1]. For this experiment, two groups of mice were established: 1) control group without MPS-cytoblockade – mice received only NPs at a dose of 300 µg; 2) group with MPS-cytoblockade – mice were administered 25 µg of 34-3C antibody and 300 µg of NPs. According to the results (Figure 1a), the application of MPS-cytoblockade increased the nanoparticle blood circulation half-life (t_1/2_) by 12.9-fold – from 0.8 ± 0.3 min to 10.3 ± 2.9 min.

**Figure 1.**
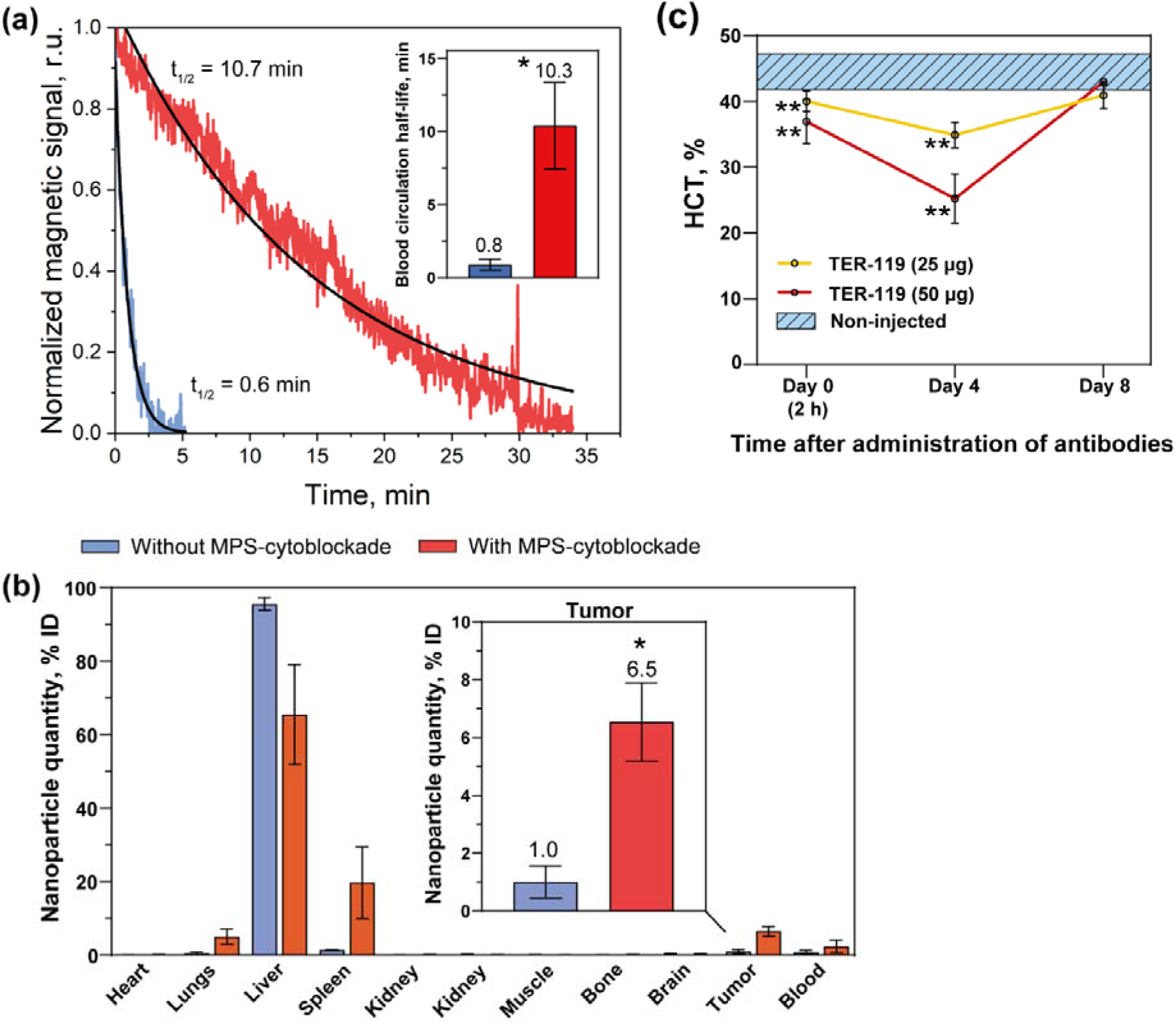
Changes in key parameters following MPS-cytoblockade. (a) Representative nanoparticle circulation kinetics and half-life times. (b) Biodistribution of nanoparticles in various organs and tissues, including the tumor. (c) Hematocrit levels on different days after administration of various doses of TER-119 antibodies to induce MPS-cytoblockade. The number of animals was n ≥ 3 for each group. Asterisks indicate significant differences: for (a) and (b) between groups with and without MPS-cytoblockade; for (c) compared to non-injected mice. The nonparametric Mann-Whitney U test was applied: * P ≤ 0.05, ** P ≤ 0.01, ns – not significant.

The model nanoparticles possessed a magnetite core and were therefore iron□containing [17]. Upon administration into an animal, the iron released from these nanoparticles can be utilized by erythroid precursors in the bone marrow, directly stimulating erythropoiesis and leading to a true increase in hematocrit – an effect not observed with nanoparticles made of inert materials. Consequently, we also investigated the effect of nanoparticle administration on changes in hematocrit levels during MPS□cytoblockade (Figure S1). The results showed that co□administration of antibodies with nanoparticles produced the same effect on hematocrit levels as antibody injection alone.

The impact of MPS-cytoblockade on the biodistribution profile and tumor accumulation of NPs was also evaluated using the MPQ method. To obtain tumor-bearing mice, 2·10^6^ CT-26 cells in 200 µL of culture medium were inoculated subcutaneously into the right flank of female BALB/c mice. Once the tumor volume reached approximately 1000 mm^3^, in a group with MPS-cytoblockade mice were administered 25 µg of 34-3C antibody 12 hours prior to injection of NPs at a dose of 1000 µg, while mice in control group without MPS-cytoblockade received only NPs at the same dose. To improve the targeting efficiency of magnetic nanoparticles to the tumor site, magnetic focusing was performed using an externally applied magnet. Following a three-hour circulation period, the animals were euthanized via cervical dislocation. Subsequently, a set of organs and tissues was harvested, including the heart, lungs, liver, spleen, kidneys, muscle, bone, brain, tumor, and blood. Each harvested sample was then transferred to the instrument’s detection coil for measurement of the magnetic signal. NP accumulation in tumor reached 1.0 ± 0.5% in control group, compared to 6.5 ± 1.3% in the MPS-cytoblockade group (Figure 1b).

The use of anti-erythrocyte antibody injection during MPS-cytoblockade and subsequent saturation of macrophages with erythrocytes is associated with a decline in hematocrit (HCT) – the ratio of red blood cell volume to total blood volume. To assess changes in this parameter with high precision, we first optimized the hematocrit study protocols.

Typically, HCT is measured to identify conditions such as anemia or polycythemia using either an automated hematology analyzer or the microhematocrit method. The microhematocrit method involves centrifuging blood samples in capillary tubes, which separates the blood into three distinct layers: red blood cells, buffy coat layer (containing white blood cells and platelets), and plasma. This method requires only a small blood volume, is simple to perform, and is inexpensive [18]. We compared these two methods for hematocrit determination. The value obtained using the hematology analyzer did not differ from that measured with the microhematocrit method (Figure S2a). Furthermore, the hematocrit levels obtained from healthy intact male BALB/c mice aligned with normal reference ranges reported in the literature [19]. Although the former method yielded a slightly higher average, this may be attributed to specific features of murine blood cell morphology. Mice have smaller erythrocyte diameters compared to other rodents, which during automated analysis on a hematology analyzer can lead to an upward shift in parameters such as red blood cell count and hematocrit [20]. For subsequent studies, we selected the microhematocrit method due to its combination of simplicity and speed of analysis.

Blood collection in laboratory mice can be performed via various methods – from retro-orbital sinus, cheek, tail vein, or heart. Since the procedure may influence measurement results due to differences in animal restraint, invasiveness, and stress, we examined its effect on hematocrit levels. Specifically, we compared values obtained by submandibular venipuncture, tail tip amputation, and intracardiac puncture against those from retro-orbital sinus collection (Figure S2b). As expected, hematocrit values did not differ significantly. For subsequent work, we selected retro-orbital sinus puncture method due to its widespread use in murine studies, relatively straightforward procedure, and ability to yield sufficient blood volumes while minimizing animal discomfort [21]. Next, using the optimized hematocrit study protocols, we investigated changes in this parameter during MPS-cytoblockade implementation.

In the original study, it was shown that the 25 µg dose of the 34-3C antibody reduces hematocrit levels by approximately 5% compared to baseline [1]. We hypothesize that administering a higher antibody dose could further prolong nanoparticle circulation time and, consequently, enhance therapeutic efficacy. Increasing the antibody dose may lead to more pronounced effects. Therefore, we investigated the dose-dependent impact of anti-red blood cell antibodies on hematocrit reduction relative to normal levels. For this purpose, TER-119 antibodies were administered intravenously via the retro-orbital sinus to male BALB/c mice at doses of 25 and 50 µg. Blood was subsequently collected from a non-injected retro-orbital sinus on days 0 (2 hours post-injection), 4, and 8. The results were compared with those from a group of non-injected control animals (Figure 1c). Both antibody doses bound to circulating erythrocytes, inducing their enhanced clearance by macrophages and leading to a corresponding decrease in hematocrit. A significant reduction was observed on day 0 and 4 post-administration. The most pronounced decline occurred on day 4: the 25 µg dose reduced hematocrit by approximately 9.5% relative to normal levels, while the 50 µg dose caused a 19.2% reduction. By day 8, however, hematocrit levels had recovered to baseline.

### 2.2. Development of a method for maintaining normal hematocrit levels in mice during MPS-cytoblockade through the administration of donor erythrocytes

The results presented in Figure 1c allowed us to estimate the required volume of donor erythrocyte suspension needed to maintain normal hematocrit levels during MPS-cytoblockade. In addition, we consulted published data: the acceptable transfusion volume of whole blood or donor erythrocyte suspension (with a hematocrit of approximately 50%) for BALB/c mice ranges from 100 to 400 µL [22], [23] and can even reach 500 µL [24]. The procedure is typically performed intravenously via the tail vein [25] or the retro-orbital sinus [26]. Based on our practical observations, however, mice tolerated the tail vein transfusion more readily, therefore, this route was selected for subsequent experiments.

Based on these data, we selected several potential transfusion protocols for the donor erythrocyte suspension. Specifically, to restore hematocrit levels following administration of the 50 µg TER-119 antibody dose, we infused mice with three different volumes (200, 300, and 400 µL) of a suspension with 50% hematocrit (Figure 2, Schemes 1-3). Prior to transfusion, the suspension was incubated with the antibodies for 5 min at room temperature and then administered via the tail vein. For the 400 µL transfusion, the volume was split into two equal parts: the first 200 µL, pre-incubated with antibodies, was administered on day 0, followed by a second 200 µL suspension on day 2. Hematocrit levels for all schemes were assessed on day 4 post-antibody injection and compared to those in non-injected control mice and mice injected with NaCl. For the schemes that optimally maintained hematocrit, we also evaluated the parameter on days 0 and 8.

**Figure 2.**
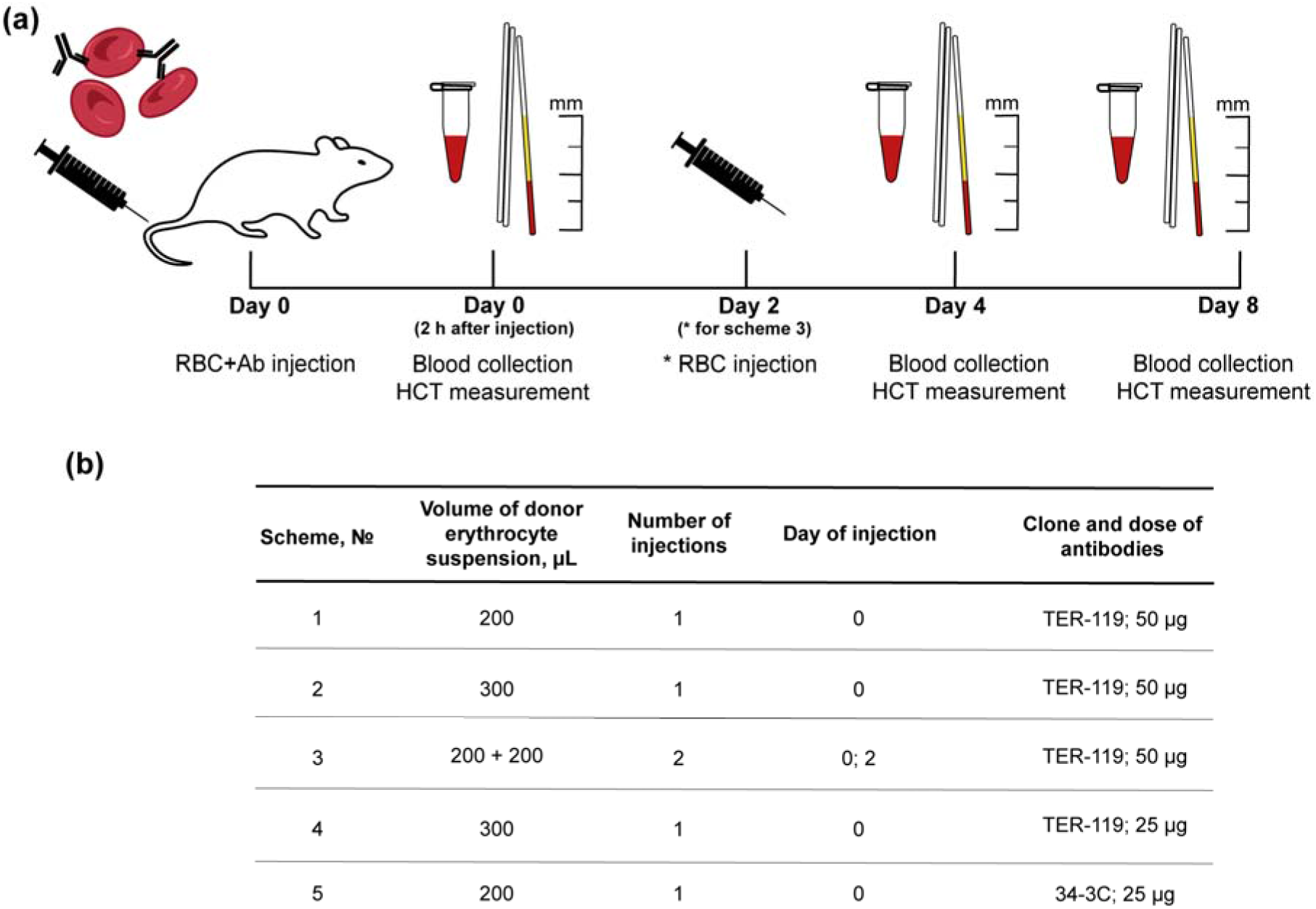
Schemes for donor erythrocyte suspension administration to maintain normal hematocrit levels in mice during MPS-cytoblockade. (a) Experimental design. (b) Summary of implemented procedures. RBC – red blood cells, Ab – antibodies.

Based on the comparison of HCT levels on day 4 after MPS-cytoblockade via injection of 50 µg TER-119 antibody (Figure 3a), Scheme 3 was the most effective at maintaining this parameter within the normal range. We therefore evaluated HCT for this scheme on day 0 (2 hours post-injection) and day 8 (Figure 3b). A slight increase was observed on day 0, likely attributable to the measurement being taken shortly after the donor erythrocyte transfusion. However, by days 4 and 8, hematocrit levels were comparable to those in non-injected control mice.

**Figure 3.**
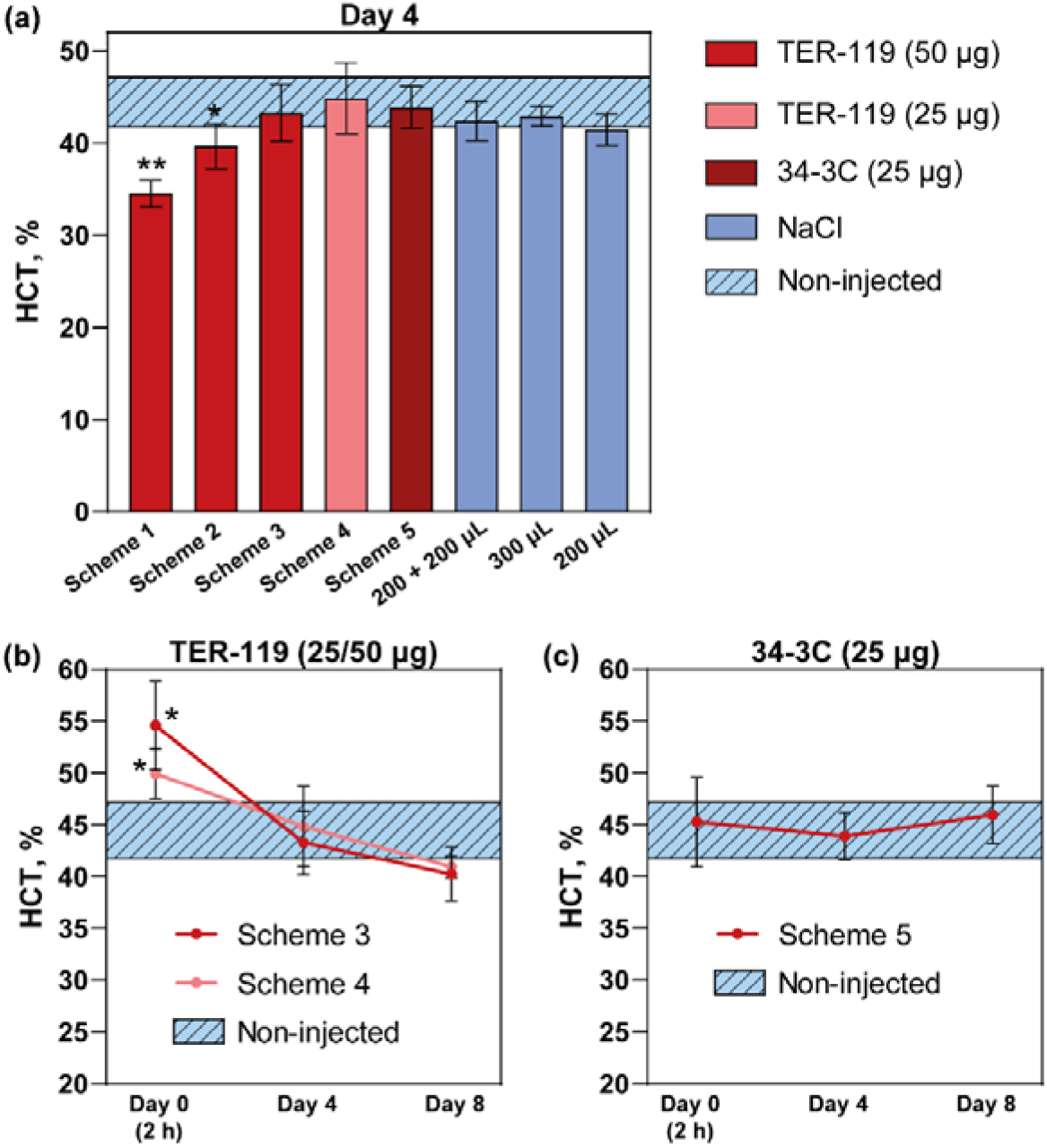
Evaluation of various protocols for hematocrit restoration using donor RBC suspension transfusion. (a) Hematocrit on day 4 after injection of TER-119 antibodies at doses of 50 μg or 25 μg, and 34-3C antibody at a dose of 25 μg, under different transfusion protocols. (b, c) Hematocrit on days 0, 4, and 8 for schemes 3, 4, and 5. The number of animals was n = 8 for the non-injected group and n ≥ 3 for the other groups. Asterisks indicate significant differences compared to non-injected mice. The nonparametric Mann-Whitney U test was applied: * P ≤ 0.05, ** P ≤ 0.01, ns – not significant.

Using the same approach, we identified optimal protocols for restoring hematocrit following administration of 25 µg of either TER-119 or 34-3C antibodies (Figure 2, Schemes 4 and 5). The proposed regimens successfully maintained normal HCT levels on days 4 and 8; Scheme 5 also sustained normal levels on day 0 post-antibody injection (Figure 3b,c).

### 2.3. Development of a method for maintaining normal hematocrit levels in mice during MPS-cytoblockade through the administration of hormone erythropoietin

The hormone erythropoietin (EPO) is clinically used to treat anemia associated with renal failure or chemotherapy for non-myeloid hematological malignancies, during preoperative erythrocyte mobilization or autologous blood donation, and in several other conditions [27]. Its mechanism of action involves binding to the EpoR receptor on erythroid progenitor cells, thereby promoting their proliferation and differentiation [28]. In the present study, EPO was administered to maintain hematocrit levels within the normal physiological range during MPS-cytoblockade by stimulating erythropoiesis.

Numerous studies have established significant homology between human EPO and the hormone from several other mammals, both in their coding DNA and amino acid sequences [29]. For instance, human EPO exhibits approximately 91% amino acid identity with simian EPO and 80-82% with porcine, ovine, murine, and rat variants. Furthermore, the biological activity of murine EPO has also been confirmed on both murine and human erythroid progenitor cells [30]. This high degree of cross-species homology in EPO structure and function allowed us to utilize recombinant human EPO (rhEPO) for injections. The rhEPO used was produced in eukaryotic cells and possesses an amino acid sequence identical to that of endogenous human EPO.

The strategy of administering rhEPO to mice has been employed in various studies [31] [32]. For instance, Mittelman et al. demonstrated that using the hormone to treat cancer-related anemia in mice with myeloma not only increased their survival but also induced tumor regression in 30-60% of the animals. However, it is crucial to apply the minimal effective dose of rhEPO, as higher doses can lead to adverse effects, such as thrombocytopenia [33] and bone loss [34].

First, we tested 10 different administration protocols for rhEPO (hereafter EPO) (Figure 4). The hormone was administered either several days before, one day before, on the same day as, or two days after the antibody administration. Hematocrit levels were then assessed on day 4 and compared with those in a non-injected control group. For the optimal protocols, hematocrit was also monitored on days 0 and 8.

**Figure 4.**
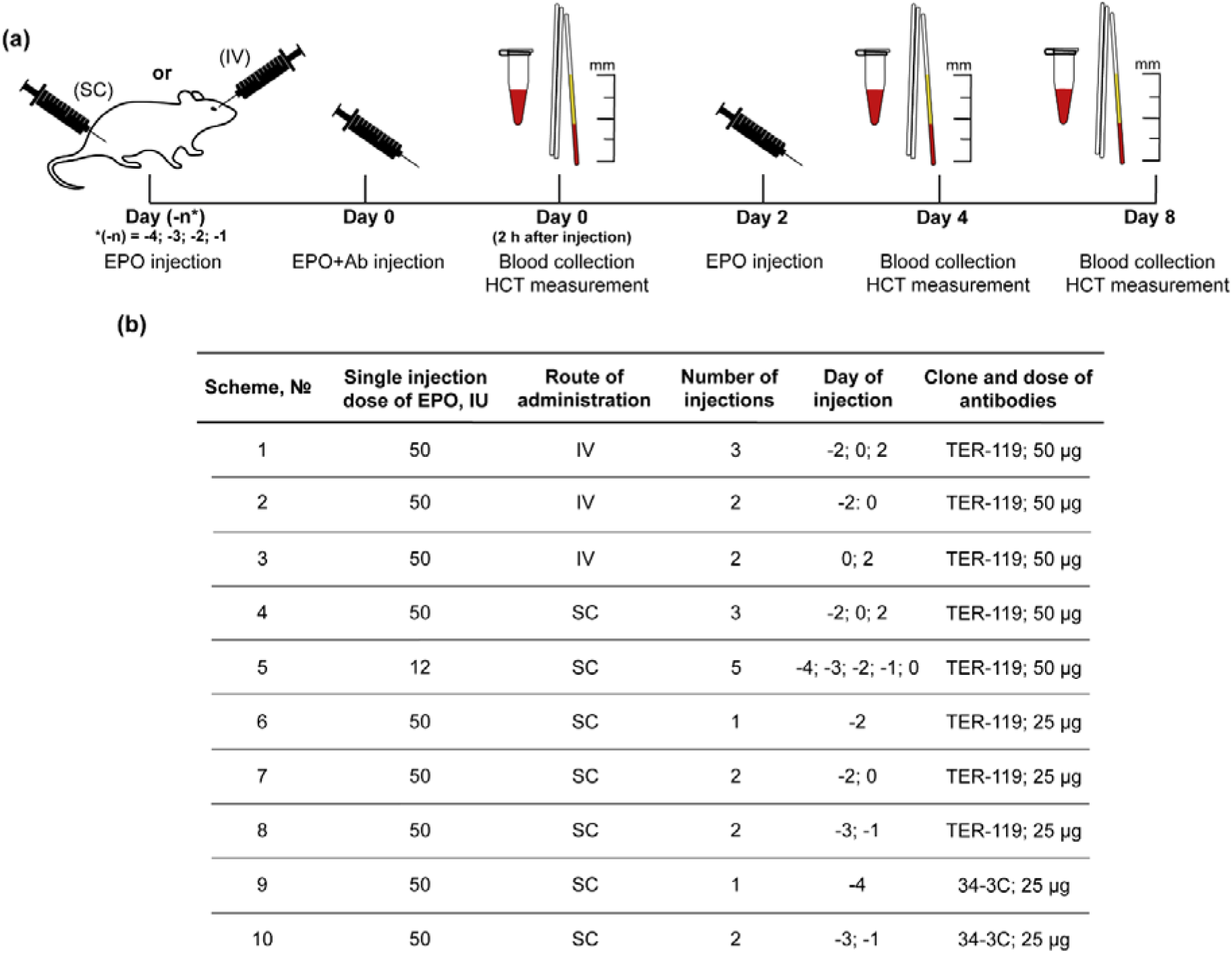
Schemes for EPO administration to maintain normal hematocrit levels in mice during MPS-cytoblockade. (a) Experimental design. (b) Summary of implemented procedures. Ab – antibodies.

To assess the dose-response relationship for EPO on hematocrit, a single injection of the hormone was administered at doses of 200, 50, 12, and 3 IU. Hematocrit values were measured on the fourth day post-injection (Figure S3). Based on the absolute mean values, the 200 IU dose increased hematocrit by 7%, 50 IU by 5%, and 12 IU by 1% relative to the non-injected controls. Furthermore, since the hormone can be administered either intravenously (IV) or subcutaneously (SC), we tested both routes during MPS-cytoblockade induced by TER-119 antibodies at a dose of 50 μg per mouse (Figure 5a). Intravenous EPO administration according to schemes 2 and 3 failed to maintain normal HCT levels, in contrast to schemes 4 and 5 with subcutaneous injection. Both intravenous and subcutaneous administration of the same EPO dose (schemes 1 and 4, respectively) maintained hematocrit within the normal range; however, due to the greater invasiveness of the IV route, SC injection was deemed preferable.

**Figure 5.**
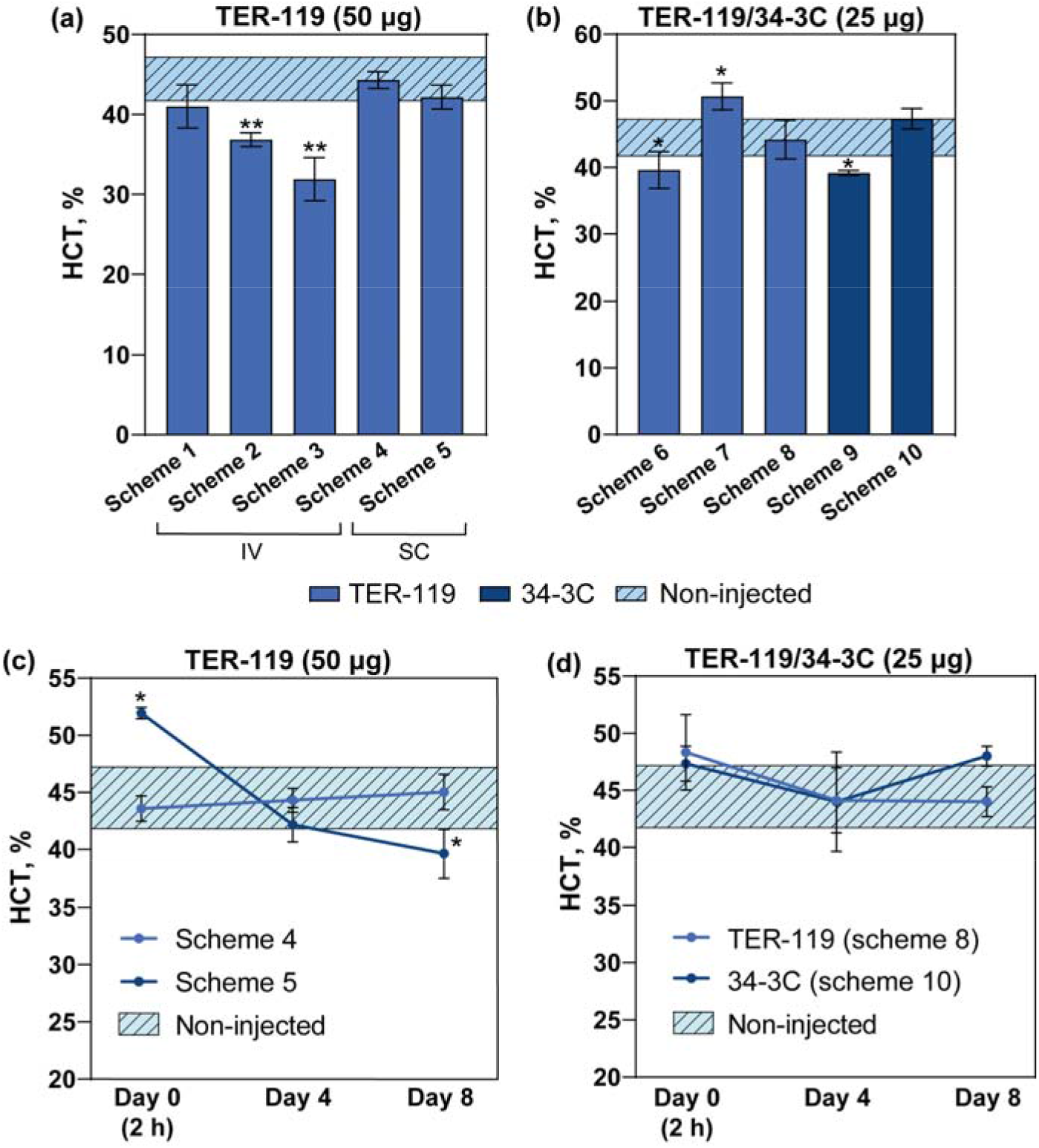
Evaluation of various protocols for hematocrit restoration using EPO injections. (a, b) Hematocrit on day 4 after administration of TER-119 (50 or 25 μg) or 34-3C (25 μg) antibodies simultaneously with EPO injections according to different schemes. (c, d) Hematocrit on days 0, 4, and 8 for schemes (c) 4, 5 and schemes (d) 8, 10. The number of animals was n = 8 for the non-injected group and n ≥ 3 for the other groups. Asterisks indicate significant differences compared to non-injected mice. The nonparametric Mann-Whitney U test was applied: * P ≤ 0.05, ** P ≤ 0.01, ns – not significant. SC – subcutaneous administration; IV – intravenous administration.

Of the two schemes (schemes 4 and 5) found to be optimal for hematocrit recovery following a 50 µg TER-119 antibody injection and showing equal efficacy on day 4, only protocol 4 sustained normal levels on both days 0 and 8 (Figure 5c). This outcome can likely be explained by the fact that in scheme 5, the five consecutive daily hormone injections before the antibody injection raised the hematocrit level too high on day 0, despite the lower total hormone dose compared to scheme 4. Furthermore, on day 8, the hematocrit level dropped significantly below normal, which could potentially be associated with more active exhaustion of erythroid progenitor cells caused by the daily injection regimen [35].

We further investigated whether normal hematocrit levels could be sustained during MPS-cytoblockade induced by lower antibody doses (25 µg) of TER-119 and 34-3С (Figure 4, schemes 6-10). For both clones, the same EPO administration protocols were effective (schemes 8 and 10): two injections – one three days before and another one day before antibody administration (Figure 5b). Using these protocols, hematocrit levels remained within the normal range on days 0, 4, and 8 (Figure 5d).

The dose of 25 µg of antibodies had already been used in previous studies [1], [13] focusing on the application of MPS-cytoblockade, while the use of higher doses, such as 50 µg, raised concerns about toxicity associated with the depletion of a large number of red blood cells. However, we demonstrated that hematocrit recovery was achieved with both antibody doses – both through the administration of donor red blood cell suspensions and through erythropoietin injections.

### 2.4. Flow cytometric analysis of reticulocyte response to different hematocrit-maintaining strategies during MPS-cytoblockade

In response to MPS-cytoblockade or during the implementation of hematocrit-maintaining strategies, the number of reticulocytes in the bloodstream may change. Reticulocytes represent the final immature stage of erythrocyte development. They are produced in the red bone marrow and, after completing their primary maturation, are released into the bloodstream, where they finally lose their residual RNA and organelles, thereby becoming mature erythrocytes. The presence of this residual RNA enables the differentiation of reticulocytes from mature erythrocytes using specialized techniques, such as flow cytometry with fluorescent dyes (e.g., acridine orange) or supravital staining methods (e.g., with brilliant cresyl blue). In this work, we employed both techniques.

Using imaging flow cytometry, we quantified the reticulocyte percentage in four groups of male BALB/c mice: 1) non-injected control mice; 2) mice receiving a single injection of 25 µg of 34-3C antibodies; 3) mice received two 50 IU subcutaneous injections of EPO administered three days and one day prior to an intravenous injection of 25 µg of the 34-3C antibody; 4) mice received a single transfusion of 150 μL of donor erythrocytes that had been pre-incubated with 25 µg of the 34-3C antibody. Blood samples were collected on the fourth day post-antibody administration and stained with Cyanine5-conjugated TER-119 antibodies for specific detection of all erythroid lineage cells. To identify the reticulocyte fraction, samples were further incubated with acridine orange (AO).

For each sample, at least 100 000 events were recorded, and a compensation matrix was applied. The gating strategy was performed in four sequential steps (Figure 6a). First, in-focus images were distinguished from out-of-focus ones by applying a threshold of Gradient Root Mean Square (RMS) > 45 in the brightfield (BF) channel. The Gradient RMS parameter serves as an indicator of image sharpness by detecting large changes of pixel values in the image [36]. Following this, a gate was applied to this focused population to select single cells by utilizing a plot of Area BF versus Aspect Ratio Intensity BF, where the Aspect Ratio Intensity was defined as the ratio of the intensity along the cell’s minor axis to that along its major axis [37]. From the identified single-cell population, we generated a fluorescence intensity histogram for the Cyanine5 (Cy5) channel to isolate the total population of erythroid blood cells. Subsequently, an AO channel fluorescence intensity histogram was constructed for these cells to quantify reticulocytes in each sample.

**Figure 6.**
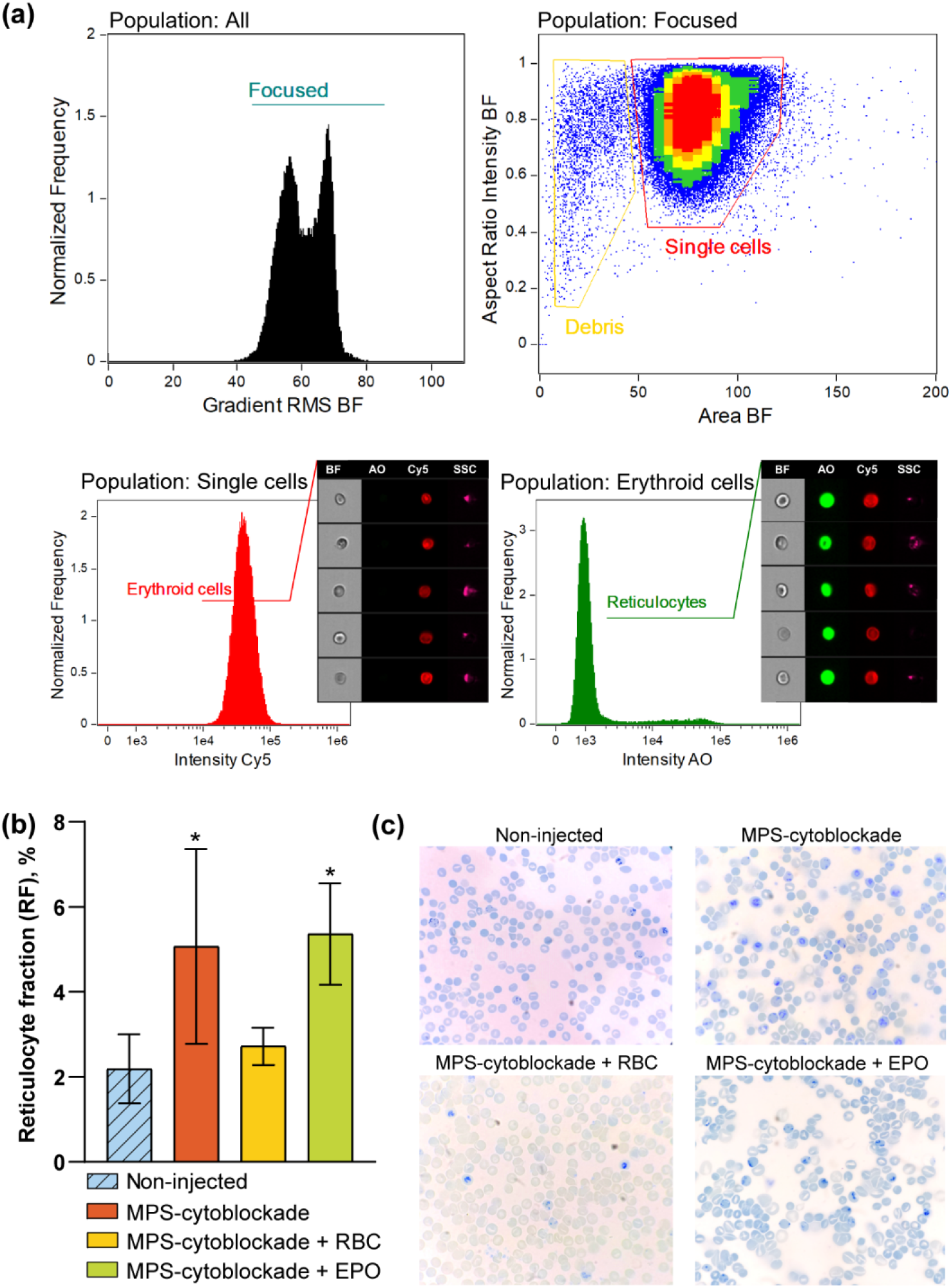
Analysis of reticulocytes using (a,b) imaging flow cytometry and (c) the supravital staining method. (a) Gating strategy illustrated for a blood sample from a mouse injected with 34-3C antibody. (b) Percentage of reticulocytes in blood samples from mice receiving a 34-3C antibody injection, a transfusion of erythrocyte suspension pre-incubated with the antibody, or EPO and antibody injections, and from non-injected control mice. (c) Representative images of blood smears stained with brilliant cresyl blue. The number of animals was n ≥ 3 for each group. Asterisks indicate significant differences compared to non-injected mice. The nonparametric Mann-Whitney U test was applied: * P ≤ 0.05, ** P ≤ 0.01, ns – not significant. Abbreviations for channels: BF – brightfield, AO – acridine orange, Cy5 – Cyanine5, SSC – side scatter.

Based on the experimental results (Figure 6b), blood samples from mice injected with the 34-3C antibody in response to induced anemia showed a significant increase in reticulocyte count (up to 5.1 ± 2.3%) compared to the control group of non-injected mice (2.2 ± 0.8%) [38,39]. A similar increase (up to 5.4 ± 1.2%) was observed in blood samples from mice receiving both EPO and antibody injections. In this case, the rise in reticulocytes could be attributed to either the hormone [40] or the antibody, however, no synergistic effect was noted, as the values obtained did not differ significantly from those in mice receiving the antibody alone. In contrast, blood samples from mice transfused with an erythrocyte suspension pre-incubated with the antibody showed no change in reticulocyte levels (2.7 ± 0.4%). This is likely because the anti-erythrocyte antibodies were almost entirely bound to the donor erythrocytes and subsequently cleared by the MPS without inducing anemia in the recipient mice.

The obtained results were further confirmed using the supravital staining method with brilliant cresyl blue. In Figure 6c reticulocytes displayed the characteristic dark blue reticular staining pattern of ribonucleic acid residues, unlike mature erythrocytes. A significant increase in reticulocyte count was observed both in blood samples from mice receiving a 34-3C antibody injection and in those receiving injections of both EPO and the antibody.

### 2.5. Investigating the influence of approaches to maintaining normal hematocrit levels on nanoparticle circulation time and tumor accumulation efficiency during MPS-cytoblockade implementation

We explored whether the proposed approaches for restoring hematocrit levels compromise the ability of MPS-cytoblockade to prolong nanoparticle circulation time. We also assessed the influence of these interventions on the NPs biodistribution profile and accumulation in the target site – the tumor.

Figure 7a shows representative nanoparticle circulation kinetic profiles. Alongside previously obtained data (as shown in Figure 1a) for groups with and without MPS-cytoblockade, this figure also includes data from mice that received MPS-cytoblockade (25 µg of 34-3C antibody and 300 µg of NPs) and were additionally treated with EPO injections (Figure 4, Scheme 10) to restore hematocrit levels. When MPS-cytoblockade was combined with EPO administration, the circulation half□life reached 14.8 ± 8.0 min, representing an 18.5□fold increase compared to the control group without MPS-cytoblockade. Importantly, both MPS-cytoblockade groups (with and without EPO injection) had t_1/2_ values that differed significantly from the group without MPS-cytoblockade (P ≤ 0.05), but did not differ from each other (ns).

**Figure 7.**
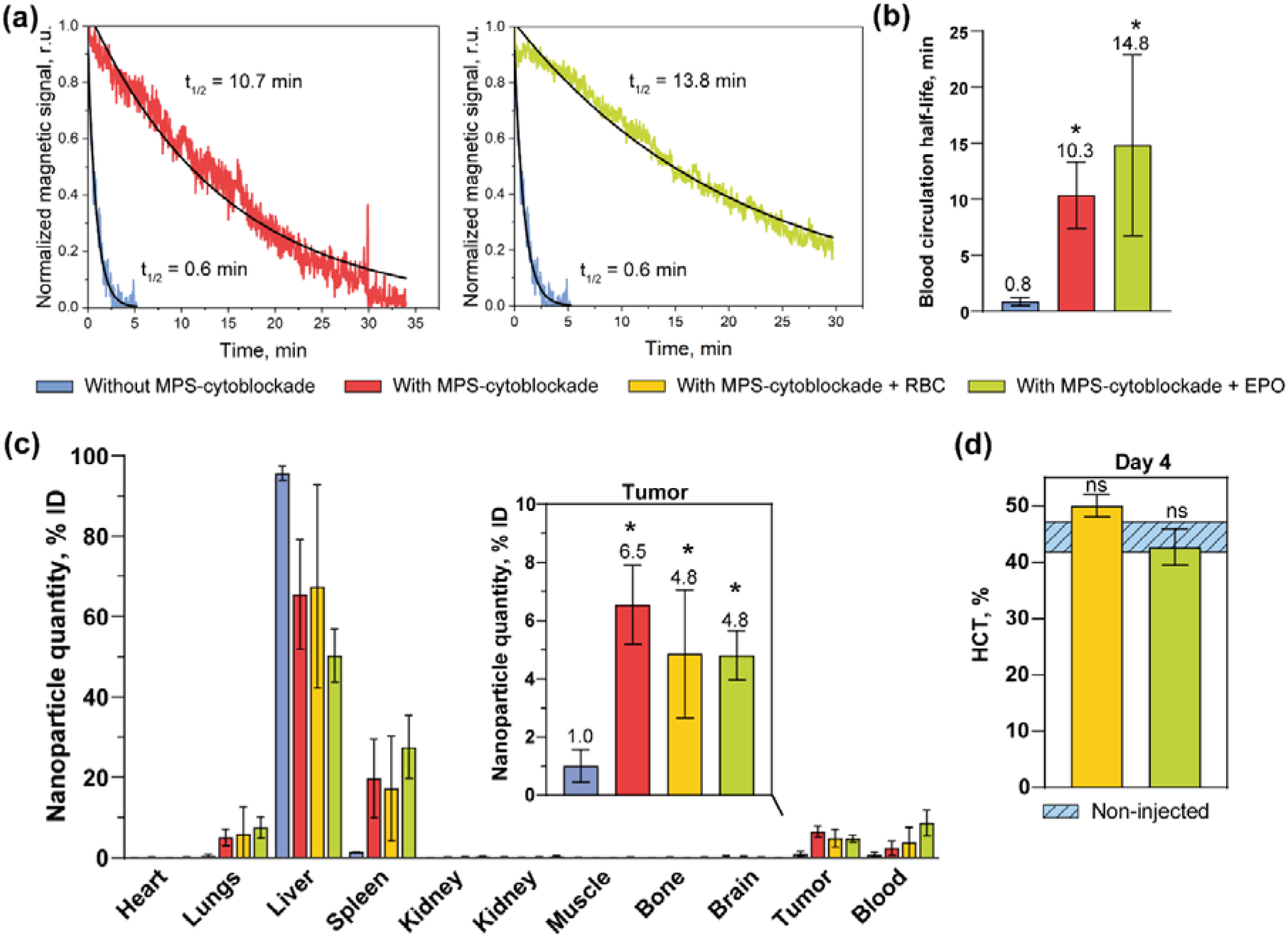
Influence of approaches to maintaining normal hematocrit levels on key parameters during MPS-cytoblockade. (a) Representative nanoparticle circulation kinetics and (b) half-life times. (c) Biodistribution of nanoparticles in various organs and tissues, including the tumor. (d) Hematocrit levels on day 4 post-MPS-cytoblockade with donor RBC suspension transfusion or EPO administration in tumor-bearing mice. The number of animals was n ≥ 3 for each group. Asterisks indicate significant differences compared to non-injected mice. The nonparametric Mann-Whitney U test was applied: * P ≤ 0.05, ns – not significant.

Similar to the data obtained earlier (Figure 1b) for groups with and without MPS-Сytoblockade, the biodistribution profile and tumor accumulation of NPs were also examined in two additional experimental groups: one receiving MPS-cytoblockade (25 µg of 34-3C antibody and 1000 µg of NPs) plus erythrocyte suspension injections (Figure 2, Scheme 5), and another receiving MPS-cytoblockade (same antibody and NP dose) plus EPO injections (Figure 4, Scheme 10), both aimed at restoring hematocrit levels. For all three MPS-cytoblockade groups – MPS-cytoblockade alone, MPS-cytoblockade with EPO or erythrocyte suspension – a similar biodistribution profile was observed. This profile was characterized by reduced liver accumulation and increased accumulation in the spleen and lungs. Moreover, nanoparticle accumulation in tumor was significantly higher compared to the control group without MPS-cytoblockade (P ≤ 0.05). No significant differences in NPs accumulation were found among the three groups, either in the tumor or in other organs/tissues.

Taken together, these results confirm that the proposed hematocrit restoration approaches do not reduce the effectiveness of MPS-cytoblockade, neither in prolonging NPs circulation time nor in enhancing NPs accumulation in target tissues.

Furthermore, we confirmed that both methods maintain hematocrit levels during MPS-cytoblockade in tumor-bearing mice. To achieve this, female BALB/c mice bearing CT-26 tumors were administered antibodies along with a donor RBC suspension according to Scheme 5 (Figure 2) or antibodies with erythropoietin according to Scheme 10 (Figure 4). Hematocrit levels were analyzed on day 4 post-antibody administration and compared to those of a non-injected control group. The results (Figure 5d) demonstrate that both hematocrit restoration protocols effectively maintained this parameter within the normal range. Therefore, the protocols optimized above in healthy mice can also be successfully applied to tumor-bearing mice.

## 3. Materials and methods

### 3.1. Materials

The following reagents were used in the experiments: Zoletil 100 (Virbac, France); Xyla (Interchemie Werken De Adelaar Eesti AS, Estonia); sodium chloride (Helicon, Russia); phosphate-buffered saline (PBS) pH 7.4 (ECO-Service, Russia); mouse anti-mouse-RBC antibody clone 34-3C (Cat. HM1120-FS) (Hycult Biotech, Netherlands); rat anti-mouse-RBC antibody clone TER-119 (Cat. BE0183) (BioXCell, USA); erythropoietin beta 2000 IU (Binnopharm, Russia); heparin 5000 IU (Belmedpreparaty, Belarus); sulfo-N-hydroxysuccinimide Cyanine5 (Lumiprobe, Russia); acridine orange (Dia-M, Russia); brilliant cresyl blue solution (Abris+, Russia); 100 nm glucuronic acid-coated nanoparticles fluidMAG-ARA (Chemicell, Germany).

### 3.2. Animals

All procedures were approved by the Institutional Animal Care and Use Committee of the Shemyakin-Ovchinnikov Institute of Bioorganic Chemistry Russian Academy of Sciences according to protocol # 375/2023 (September 20, 2023–September 19, 2026). Male (22-24 g, 2-4 months) and female (18-22 g, 2-3 months) BALB/c mice were used. For various manipulations, the animals were anesthetized intraperitoneally with a combination of Zoletil 100/Xyla at a dose of 40/1.5 mg/kg. To obtain tumor-bearing mice, 2×10^6^ CT26 cells in 200 µL of serum-free DMEM medium were injected subcutaneously into the right flank of the female mice.

### 3.3. Measuring hematocrit level

Mice were anesthetized, and blood was collected from the retro-orbital sinus (unless otherwise stated) into a tube containing EDTA K^+^ and mixed thoroughly. Using an automatic pipette, 60 µL of blood was collected and placed in a microhematocrit capillary tubes, which were centrifuged at 4800 rpm for 16 min in an SH120-1S centrifuge (Armed, Russia). Hematocrit was determined by measuring the proportion of red blood cells relative to the total blood volume, expressed as a percentage. We also used the automated hematology veterinary analyzer 5180 Vet (URIT Medical Electronic Co., Ltd., China) to determine this parameter. The test tube with 40 µL of blood was placed under the needle of the analyzer sampler. Three measurements were taken for each sample.

### 3.4. Obtaining a donor red blood cell suspension for the transfusion procedure

For subsequent blood collection, 100 μL of heparin was administered intravenously via the retro-orbital sinus to male BALB/c mice. Blood was then collected from the other retro-orbital sinus. The obtained blood was centrifuged for 15 min at 1500 g, after which the erythrocytes were separated from the plasma. Erythrocytes were washed three times in sterile PBS by centrifugation and finally resuspended to a 50% hematocrit in the same solution.

### 3.5. Transfusion of donor red blood cell suspension and erythropoietin administration

To visualize the lateral tail vein, the tail of a male BALB/c mouse was warmed by immersion in 40°C water. A 0.5 mL insulin syringe with a 29G needle was then inserted into the middle third of the vein, holding the syringe parallel to the tail. After injection, the needle was gently withdrawn, and a gauze pad was held firmly against the injection site for 10-15 seconds to prevent bleeding.

Erythropoietin was administered subcutaneously or intravenously to male BALB/c mice (unless otherwise indicated) into the right hind thigh or retro-orbital sinus, respectively, using a 1-mL insulin syringe with a 29G needle.

### 3.6. Conjugation of TER-119 anti-erythrocyte antibodies with a fluorescent label

TER-119 antibodies were conjugated to a sulfo-Cyanine5 NHS ester dye (Cy5). For this, 25 µL of antibodies (7.56 g/L) were mixed with 17.5 µL of the fluorescent dye (1 g/L) and incubated at 37 °C for two hours. The resulting conjugate was subsequently purified from unbound dye using a Zeba Spin Desalting Column 7K MWCO (Thermo Fisher Scientific, USA) according to the manufacturer’s protocol.

### 3.7. Analysis of reticulocytes using imaging flow cytometry

Whole blood was collected from the retro-orbital sinus of male BALB/c mice into EDTA K^+^ tubes. For staining, 5 µL of blood was diluted in 1 mL of PBS. From this dilution, two 200 µL aliquots were prepared: 1) an unstained control for autofluorescence measurement; 2) a sample stained with 2 µL of acridine orange (AO, 5 mg/L) and 1 µL of Cy5-conjugated TER-119 antibodies at a concentration of 7.5 g/L. Both aliquots were incubated for 30 min at room temperature, protected from light. After incubation, the cells were washed with PBS and analyzed on an ImageStreamX Mk II imaging flow cytometer (Cytek Biosciences, USA). A total of 100 000 events were acquired for each sample. A detailed description of the gating procedure is provided in the Results and Discussion section (Figure 4a).

Single-stained controls for compensation were prepared and acquired separately. Whole blood samples were prepared: 1) unstained control; 2) stained with Cy5-conjugated TER-119 antibodies only; 3) stained with AO only. A total of 1000 events were recorded for each sample. The compensation matrix was calculated, and subsequent data analysis was performed using the IDEAS analysis software (Cytek Biosciences, USA).

### 3.8. Analysis of reticulocytes using the supravital staining method

Whole blood was collected from the retro-orbital sinus of male mice into a tube containing heparin and mixed gently. A 100 µL aliquot of blood was combined with an equal volume of brilliant cresyl blue solution (ABRIS+, Russia) and incubated for 20 min at room temperature. Subsequently, 10 µL of the mixture was used to prepare a blood smear on a glass slide. The smear was examined under an Olympus CX23 light microscope (Olympus, Japan).

### 3.9. Measurement of nanoparticle circulation kinetics in the bloodstream of mice

The *in vivo* circulation kinetics of nanoparticles were characterized using the MPQ method. For the measurements, the tail of an anesthetized male mouse was positioned within the instrument’s detection coil. Subsequently, a single dose of magnetic nanoparticles (300 µg in 100 µL of a 5% glucose solution) was administered via intravenous injection into the retro-orbital sinus. The magnetic signal corresponding to circulating nanoparticles was monitored continuously, with data points acquired every 1.6 seconds until the signal intensity diminished entirely to the baseline noise level. To calculate the circulation half-life of the nanoparticles, the recorded kinetic data were normalized relative to the peak signal intensity and plotted as a function of time. The segment of the decay curve representing a drop in the normalized signal from 0.9 to 0.1 was fitted via an exponential model: y = a*e^(bx). Then the circulation half-life was derived using the equation: t_1/2_ = ln(2)/(–b).

### 3.10. Study of nanoparticle biodistribution

The biodistribution profile of the nanoparticles was evaluated using the MPQ method. Mice received a single intravenous dose of magnetic nanoparticles (1000 µg in 100 µL of a 5% glucose solution) via the retro-orbital sinus. To improve the targeting efficiency to the tumor site, magnetic focusing was performed using an externally applied neodymium magnet (20 × 10 × 5 mm). Following a 3-hour circulation period, the animals were euthanized. Organ or tissue samples (∼ 50 mg each) were subsequently harvested from the heart, lungs, liver, spleen, kidneys, muscle, bone, brain, tumor, and blood. To quantify the relative nanoparticle accumulation in each organ or tissue, the integral magnetic signal from that sample was divided by the sum of the integral signals from all collected organs and tissues.

### 3.11. Statistical analysis

All studies were performed with group sizes (n) specified in the corresponding figure legends. Data are presented as mean ± standard deviation. Comparisons between two groups were analyzed using a nonparametric Mann-Whitney U test. Statistical significance was defined as follows: ns (not significant) for P > 0.05; * for P ≤ 0.05; and ** for P ≤ 0.01.

## 4. Conclusions

A range of strategies extends beyond MPS-cytoblockade to address the rapid systemic clearance of nanomaterials from the bloodstream. Established approaches, however, are constrained by significant drawbacks. Surface engineering with stealth coatings (e.g., polyethylene glycol [41]) risks inducing an immune response and can interfere with the functional architecture of sophisticated nanocarriers [6]. Alternative methods that directly modulate the MPS, such as depletion of phagocytic cells with clodronate [42], incur irreversible toxicity [8,43], while complex cell-hitchhiking techniques lack universality and require non-trivial preparation [44,45].

MPS-cytoblockade circumvents these limitations: it obviates the need for nanomaterial redesign, exerts a comparatively mild inhibitory effect on the MPS [13], and demonstrates universal compatibility with diverse nanoagents. This technique exhibits profound efficacy, enhancing the circulatory half-life of nanoparticles by up to 32-fold [1]. Nevertheless, its implementation induces a transient reduction in hematocrit, a physiological side effect that constitutes the principal translational challenge for this otherwise promising platform. This effect is particularly critical in oncology, where even a transient drop in hematocrit can exacerbate the pre-existing anemia commonly seen in cancer patients and potentially worsen their clinical status [15].

Here, we demonstrate that the hematocrit reduction associated with the MPS-cytoblockade can be effectively managed. We have developed and validated protocols to restore hematocrit levels following the administration of various doses and clones of anti-erythrocyte antibodies. This was achieved via two distinct strategies: 1) the infusion of a donor erythrocyte suspension pre-incubated with the anti-erythrocyte antibodies, or 2) the administration of the hormone erythropoietin. The first strategy relies on the preferential binding of antibodies to the infused donor erythrocytes, thereby directing the enhanced clearance primarily toward the donor cells and exerting minimal impact on the recipient’s own RBCs. The second strategy involves inducing erythropoiesis directly by stimulating RBC production in the bone marrow. Critically, both interventions preserve the core therapeutic benefit of the MPS-cytoblockade – a dramatically extended nanoparticle circulation time. Furthermore, they do not alter the biodistribution of the nanoagents, ensuring uncompromised tumor delivery efficiency.

Thus, the results presented in this work provide a crucial safety optimization for MPS-cytoblockade. By removing this major physiological barrier, we advance the technology’s clinical translation and bring targeted nanotherapeutics a step closer to realizing their full potential.

## Author Contributions

Elizaveta N. Mochalova: conceptualization, methodology, validation, formal analysis, investigation, writing - original draft, writing - review & editing, supervision, project administration, funding acquisition; Maria A. Yurchenko: methodology, validation, formal analysis, investigation, data curation, writing - original draft, writing - review & editing, visualization; Marina P. Timofeeva: investigation; Darina A. Maedi: investigation; Petr I. Nikitin: methodology; Maxim P. Nikitin: conceptualization, resources, funding acquisition.

## Acknowledgments

We thank Maria I. Ilyina for conducting the experiment to determine the number of reticulocytes in blood samples using the supravital staining method. Different parts of this multidisciplinary study were supported by the Russian Science Foundation project № 24-74-00040.

## Declaration of generative AI and AI-assisted technologies in the writing process

During the preparation of this work the authors used DeepSeek Chat in order to improve language. After using this service, the authors reviewed and edited the content as needed and take full responsibility for the content of the publication.

## Data availability statement

The data supporting this study are available from the corresponding author upon request.

