## Supplementary materials for "Development of approaches to overcome the drop in hematocrit when implementing mononuclear phagocyte system cytoblockade *in vivo* used to prolong the circulation of nanoparticles in the blood"

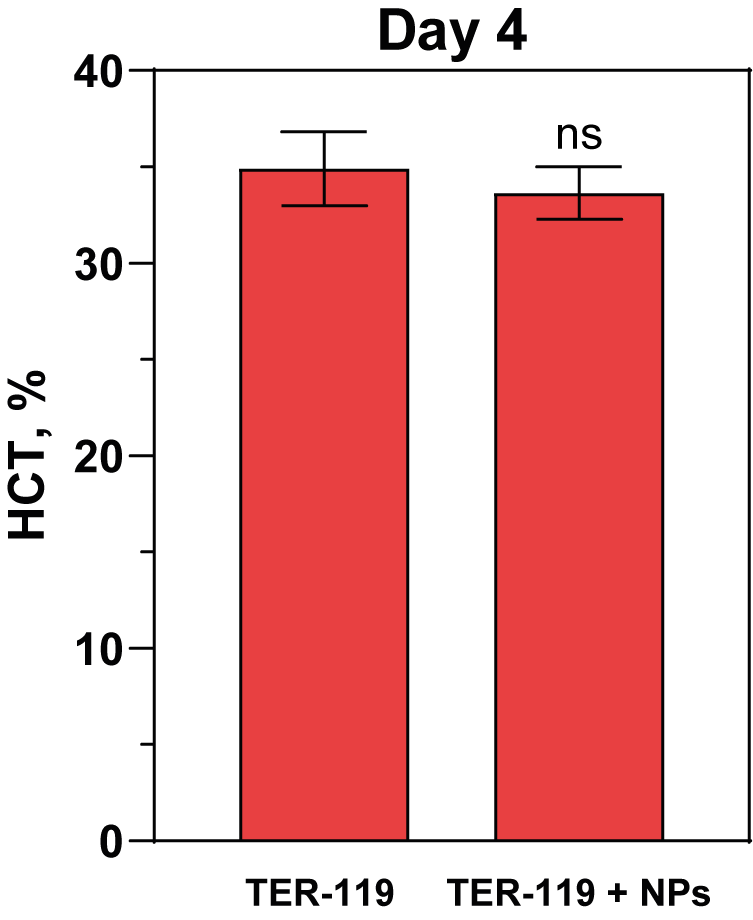

Figure S1. Effect of nanoparticle administration on changes in hematocrit (HCT) levels during MPS‑cytoblockade. MPS‑cytoblockade was induced using TER‑119 antibodies at a dose of 25 μg; nanoparticles were administered at a dose of 300 μg. Hematocrit levels were measured on day 4 after antibody injection. The number of animals was n ≥ 3 for each group. The nonparametric Mann-Whitney U test was used to assess significant differences relative to the group injected with antibodies alone: ns – not significant.

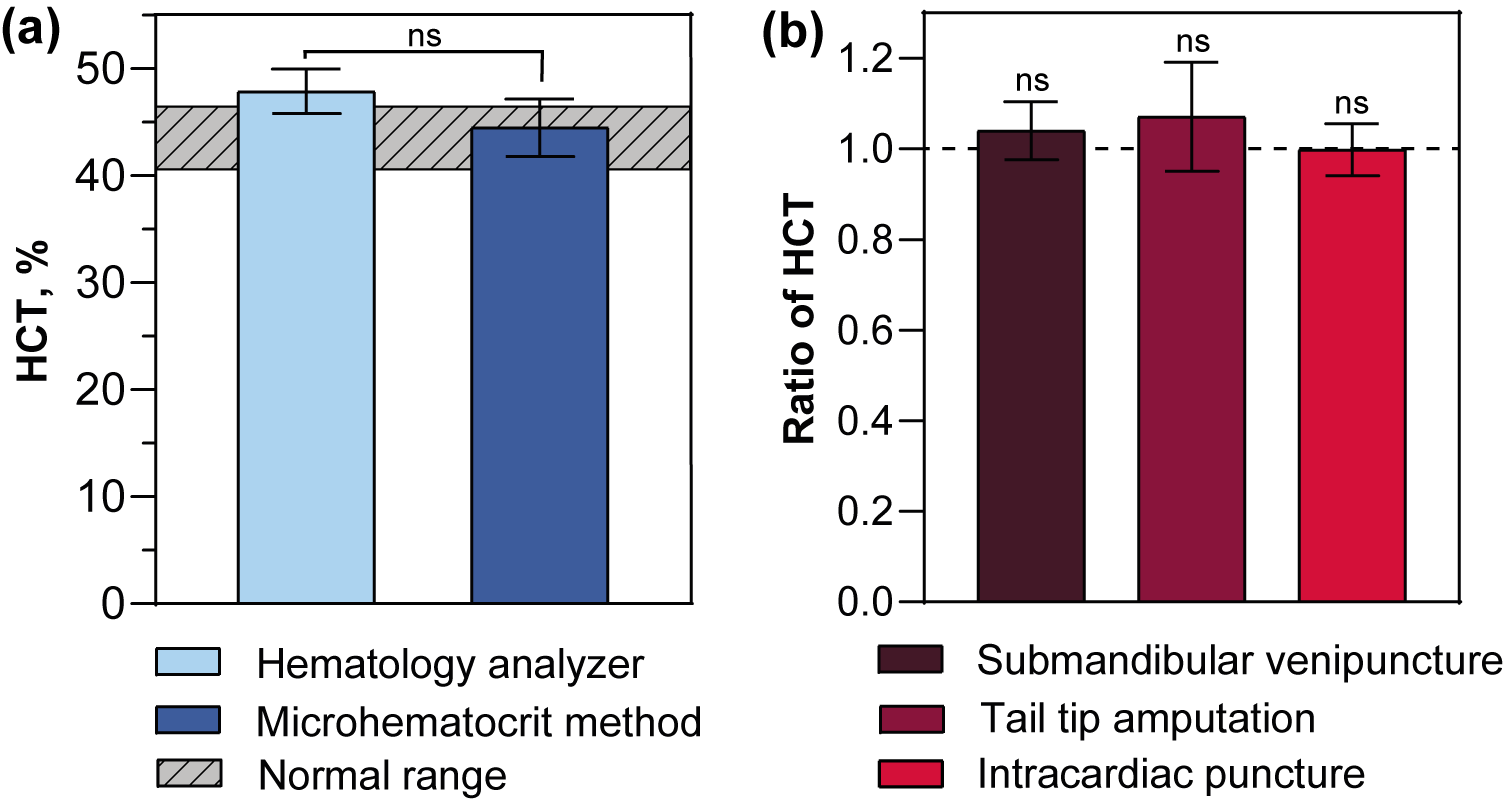

Figure S2. Optimization of hematocrit study protocols. (a) Hematocrit values obtained using a hematology analyzer and microhematocrit method; the normal range is described in the literature [1]. (b) Relative hematocrit values for various blood collection techniques normalized to retro-orbital sinus sampling. The number of animals was n ≥ 3 for each group. The nonparametric Mann-Whitney U test was used to assess significant differences (a) between the two methods, and (b) between each method and the retro-orbital sinus puncture group: ns – not significant.

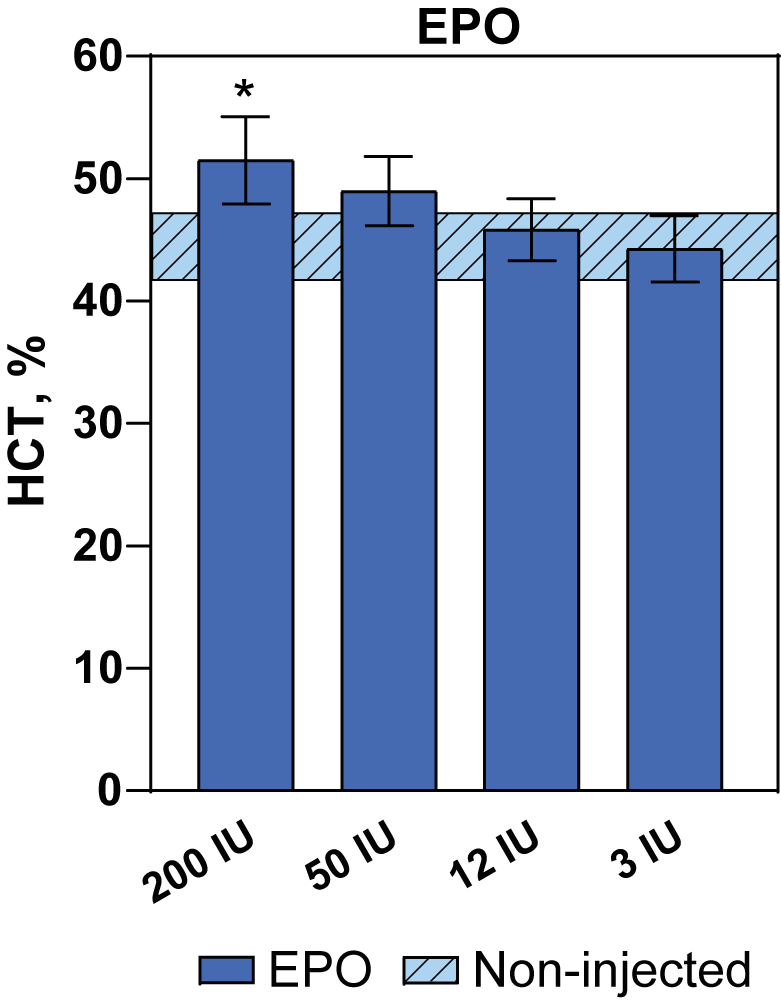

Figure S3. Hematocrit on day 4 after injection of erythropoietin (EPO) at different doses.
